# PIWIL4 regulates gene expression and piRNA levels in RSV-infected airway epithelial cells

**DOI:** 10.64898/2025.12.05.692659

**Authors:** Tiziana Corsello, Tianshuang Liu, Andrzej S Kudlicki, Yuanyi Zhang, Nicholas Dillman, Sehar Fazal, Roberto P Garofalo, Antonella Casola

## Abstract

Respiratory syncytial virus (RSV) is a leading cause of acute lower respiratory tract infections with significant morbidity and mortality in young children, the elderly and immunocompromised hosts. Despite its clinical burden, no effective RSV vaccine or therapy exists for infants, only prophylactic treatment. Small non-coding RNAs have emerged as important regulators of host-pathogen interactions. PIWI-interacting RNAs (piRNAs) are a distinct class of small non-coding RNAs known for maintaining the genome complexity and integrity in gonadal cells. However, there is growing evidence of their role in controlling gene expression in somatic cells. The biogenesis and function of piRNAs is associated with P-element Induced Wimpy testis (Piwi) proteins, whose function in the respiratory epithelium in response to infections remains largely unexplored. Here, we characterize the expression and function of the Piwi-like protein PIWIL4 in the context of RSV infection. We found that PIWIL4 is expressed in both primary and immortalized small airway epithelial cells and is significantly induced at the mRNA and protein levels following RSV infection or poly I:C stimulation, a proxy of viral infection. Immunofluorescence microscopy revealed that PIWIL4 was primarily nuclear in uninfected cells but translocated to the cytoplasm upon RSV exposure. While siRNA-mediated knockdown of PIWIL4 did not significantly affect RSV replication, it led to decreased secretion of several cytokines, chemokines and growth factors, indicating a role in modulating host innate immune responses. Transcriptomic analysis of PIWIL4-silenced iSAE cells showed significant changes in gene expression both in basal conditions and upon RSV infection. Ingenuity Pathway analysis of differentially expressed genes underscored the role of PIWIL4 in modulation of interferon signaling, cytokine production, stress and metabolic responses, as well as airway remodeling pathways. Silencing of PIWIL4 also resulted in global alteration of piRNA expression both in uninfected and infected cells. However, the predicted targets of the differentially expressed piRNAs had limited overlap with the differentially expressed genes identified by transcriptomics, suggesting a function of PIWIL4 in regulating airway epithelial cell responses at least in part independent of piRNAs. Taken together, our study uncovers an important role for PIWIL4 in somatic cells and position it as a key regulator of airway epithelial innate immunity. A better understanding of the mechanisms by which PIWIL4 affects host cells responses following a pathogen exposure may identify novel therapeutic strategies for RSV, as well as other viral respiratory infections.

## 1. Introduction

Acute lower respiratory tract infections (LRTIs) remain a leading cause of morbidity and mortality in infants and young children worldwide, with respiratory syncytial virus (RSV) as the predominant viral pathogen^1^, responsible for over 100,000 deaths with most deaths coming from low- or middle-income countries^2^. In addition to acute morbidity, RSV infection has been linked to both the development and the severity of asthma and long-term pulmonary complications^3^. No effective treatment or vaccine yet exists for RSV in children, although immunoprophylaxis is available, and immunity is incomplete, resulting in recurrent infections throughout life (8).

The P-element Induced Wimpy Testis (PIWI) family of proteins is a subclass of the Argonaute protein family, originally identified for its essential role in germline development^4^ and transposon silencing in Drosophila melanogaster^5^. PIWI protein family consists of four members PIWIL1, PIWIL2, PIWIL3, and PIWIL4, in mammalian cells ^6^. In addition to gametocytes, they have been detected in somatic tissues^7^ and implicated in a broad range of cellular processes, including gene regulation, genome stability, stem cell maintenance ^8^, and oncogenesis ^9^. A known mechanism of PIWI protein function involves their interaction with PIWI-interacting RNAs (piRNAs), a distinct class of small non-coding RNAs approximately 24–31 nucleotides in length. Together, they form piRNA-induced silencing complexes (piRISCs) that can regulate gene expression through both post-transcriptional and epigenetic mechanisms ^10^. While this pathway has been well characterized in the germline, the functional relevance of piRNAs and PIWI proteins in somatic cells is still under investigation. Emerging evidence suggests that PIWI proteins may participate in host-pathogen interactions. Recent studies in a mouse model of viral and bacterial infection found for the first time an important role for MIWI2, the mouse homologue of the human PIWIL4, in pulmonary innate immune response and disease pathogenesis^11, 12^. However, still little is known regarding the function of PIWI proteins in human airway models of infections.

Recent studies in our laboratory found a time-dependent increase in piRNA expression in RSV-infected primary airway epithelial cells ^13^ and identified piRNAs as a significant component of biologically active exosomes released from RSV-infected airway epithelial cells ^14^. This study was designed to investigate the expression and function of PIWIL4 in human airway epithelial cells in response to RSV infection. We demonstrated that PIWIL4 was inducible by RSV and poly I:C stimulation and translocated from the nucleus to the cytoplasm upon RSV infection. Using transcriptomic and piRNA profiling approaches, we showed that PIWIL4 knockdown significantly altered gene and piRNA expression levels both in basal condition and in response to infection, revealing an important role for the PIWI/piRNA axis in viral-induced airway epithelial cell responses. As the predicted targets of the differentially expressed piRNAs had limited overlap with the differentially expressed genes identified by transcriptomic analysis, our findings suggest a piRNA-independent function of PIWIL4 in regulating these responses. Our study position PIWIL4 as a previously unrecognized regulator of viral-induced human airway epithelial cell responses, opening new avenues for future therapeutic intervention in the context of respiratory infections.

## 2. Results

### 2.1 Changes in PIWIL4 expression and cellular localization in airway epithelial cells following RSV infection

Although PIWI proteins were originally considered germline restricted^4^, they can be expressed in somatic cells from different tissues/organs ^8, 15^, with higher levels of some of these proteins in specific type of cancer cells^16^. As RSV infection significantly changes piRNA expression in airway epithelial cells, we aimed to investigate the potential role of PIWI proteins in cellular responses. Based on literature reports, we decide to examine PIWIL2 and PIWIL4, two of the four PIWI isoforms, which show significant expression during normal lung development^17^, compared to PIWIL1 and PIWIL3 which are highly expressed in lung cancer cell lines^18, 19^.

In initial studies, PIWIL2 showed low detectable expression and no induction in our in vitro experimental model of RSV infected primary human small airway epithelial (SAE) cells (data not shown). To determine whether PIWIL4 expression was affected in response to a viral exposure, SAE were infected with RSV or stimulated with poly I:C, a synthetic analog of double-stranded RNA that mimics innate immune sensing of viral RNA. Cells were harvested at 6-, 15-, and 24-hours after RSV or Poly I:C treatment and total RNA and proteins were extracted for RT-qPCR and Western blot analysis. Both PIWIL4 mRNA and protein were detectable in basal conditions, and there was a progressive, time-dependent increase in its expression upon RSV infection and Poly I:C stimulation (**Figure 1**). PIWIL4 mRNA levels increased at 15 hours p.i and remained stable at 24 hours p.i. in response to RSV, while poly I:C treatment showed a more robust and progressive increase starting at 15 hours and continuing up to 24 hours post-exposure (**Figure 1A**). These findings were corroborated at the protein level by Western blot analysis, with a pattern of PIWIL4 protein induction following RSV infection and poly I:C stimulation similar to the one observed for gene expression (**Figures 1B**). A similar trend was observed in immortalized SAE (iSAE) cells, with RSV and poly I:C treatment both inducing robust PIWIL4 mRNA (**Figure 2A**) and protein expression (**Figure 2B**), starting at 15 hours and peaking at 24 hours post-exposure. Based on our initial results and previously published works, we proceeded with using iSAE cells, which exhibit behavior comparable to primary SAE cells ^20–24^, for all subsequent experiments.

**Figure 1.**
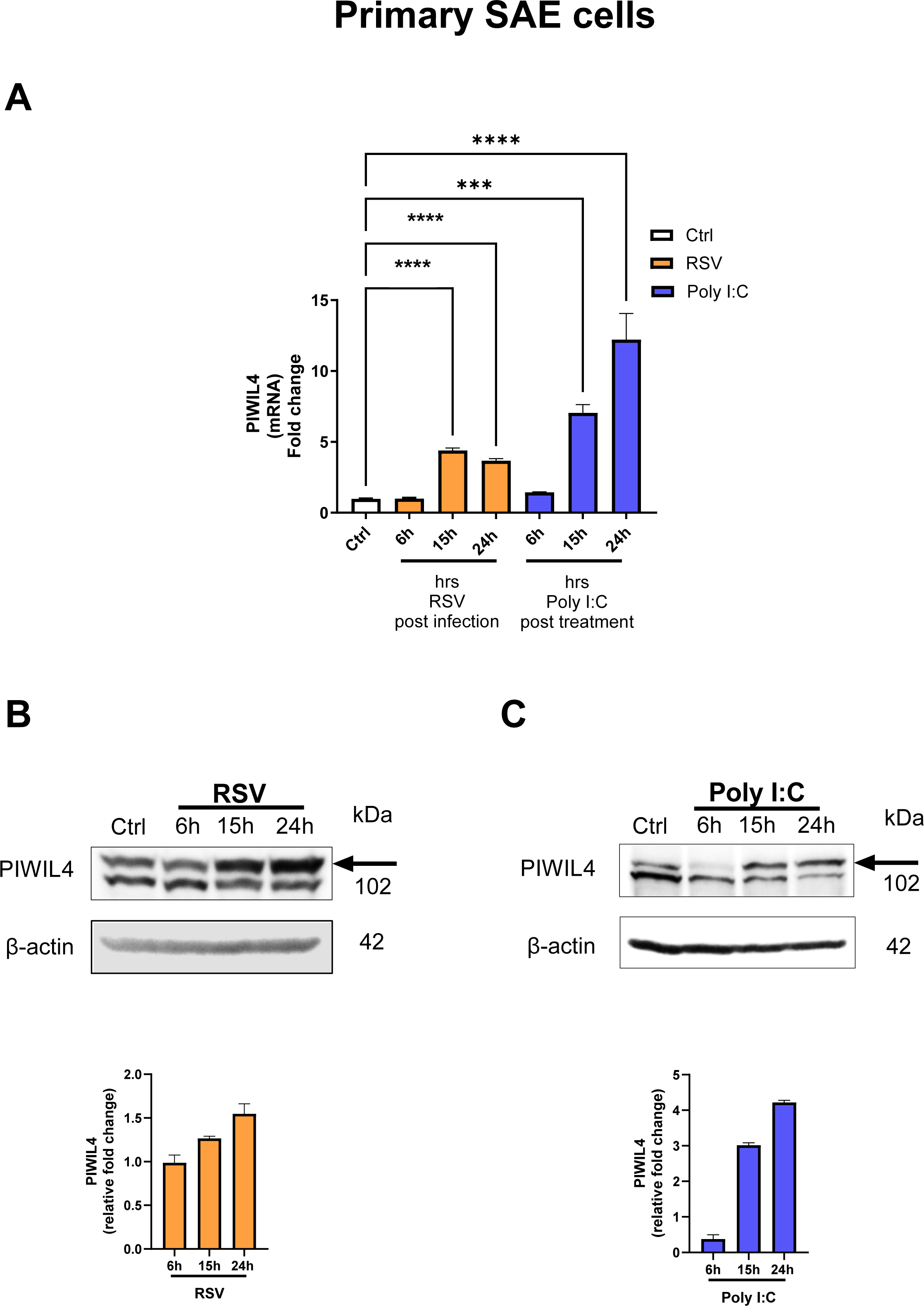
PIWIL4 mRNA and protein expression in primary small airway epithelial (SAE) cells following RSV infection and Poly I:C treatment. SAE cells control (Ctrl), infected with RSV or treated with Poly I:C were harvested at 6, 15, and 24 hours to either extract total RNA or prepare total cell lysates. (A) PIWIL4 mRNA levels measured by RT-qPCR. Data are means ± SEM (n=3, from three independent experiments run in triplicate); ***p<0.001, ****p<0.0001 infected/treated vs. control by one-way ANOVA followed by Tukey’s multiple comparisons test. (B) PIWIL4 protein levels in SAE cells following RSV infection or (C) Poly I:C treatment were measured by western blot analysis using an antibody anti-PIWIL4. Membranes were stripped and re-probed with anti-β-actin for loading control. Images are representative of two independent experiments. Graphs show relative protein fold change determined by quantification of bands and normalization to the loading control (β-actin) expressed as mean ± SEM.

**Figure 2.**
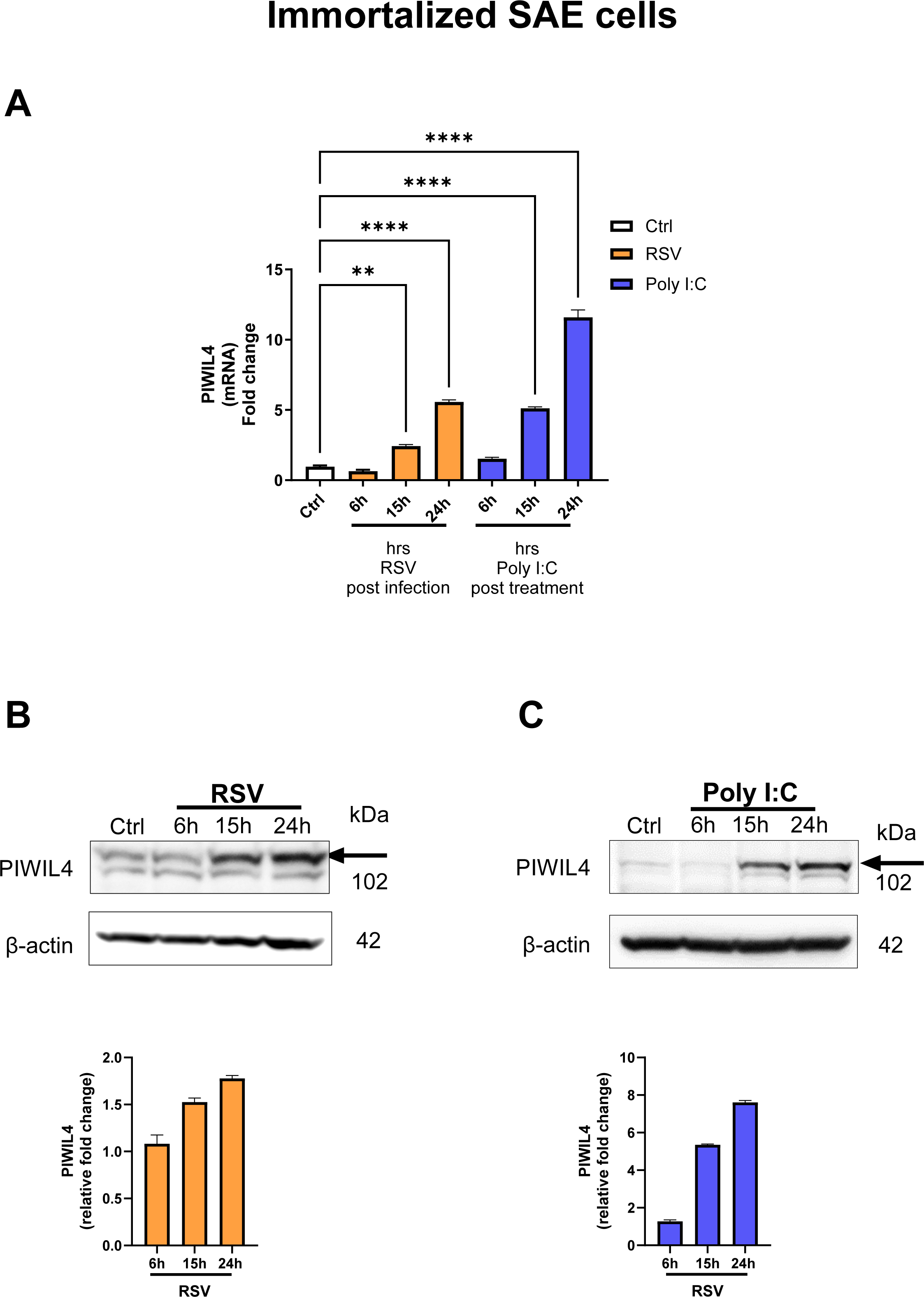
PIWIL4 mRNA and protein expression in immortalized small airway epithelial (iSAE) cells following RSV infection and Poly I:C treatment. **(A)** iSAE cells control (Ctrl), infected with RSV or treated with Poly I:C were harvested at 6, 15, and 24 hours to either extract total RNA or prepare total cell lysates. **(A)** PIWIL4 mRNA levels measured by RT-qPCR. Data are means ± SEM (n=3, from three independent experiments run in triplicate); **p<0.01, ***p<0.001, ****p<0.0001 infected/treated vs. control by one-way ANOVA followed by Tukey’s multiple comparisons test. **(B)** PIWIL4 protein levels in iSAE cells following RSV infection or **(C)** Poly I:C treatment were measured by western blot analysis using an antibody anti-PIWIL4. Membranes were stripped and re-probed with anti-β-actin for loading control. Images are representative of two independent experiments. Graphs show relative protein fold change determined by quantification of bands and normalization to the loading control (β-actin) expressed as mean ± SEM.

To explore the cellular distribution of PIWIL4, we performed immunofluorescence staining in RSV-infected and uninfected cells. In uninfected (Ctrl) cells, PIWIL4, green fluorescence, was predominantly localized to the nucleus; however, RSV infection triggered a marked redistribution of PIWIL4 from the nucleus to the cytoplasm by 15 hours p.i. (**Figure 3**). DAPI, blue fluorescence, and an antibody anti-G3BP, a marker for stress granules, red fluorescence, were used to counterstain nucleus and cytoplasmic compartments, respectively.

**Figure 3.**
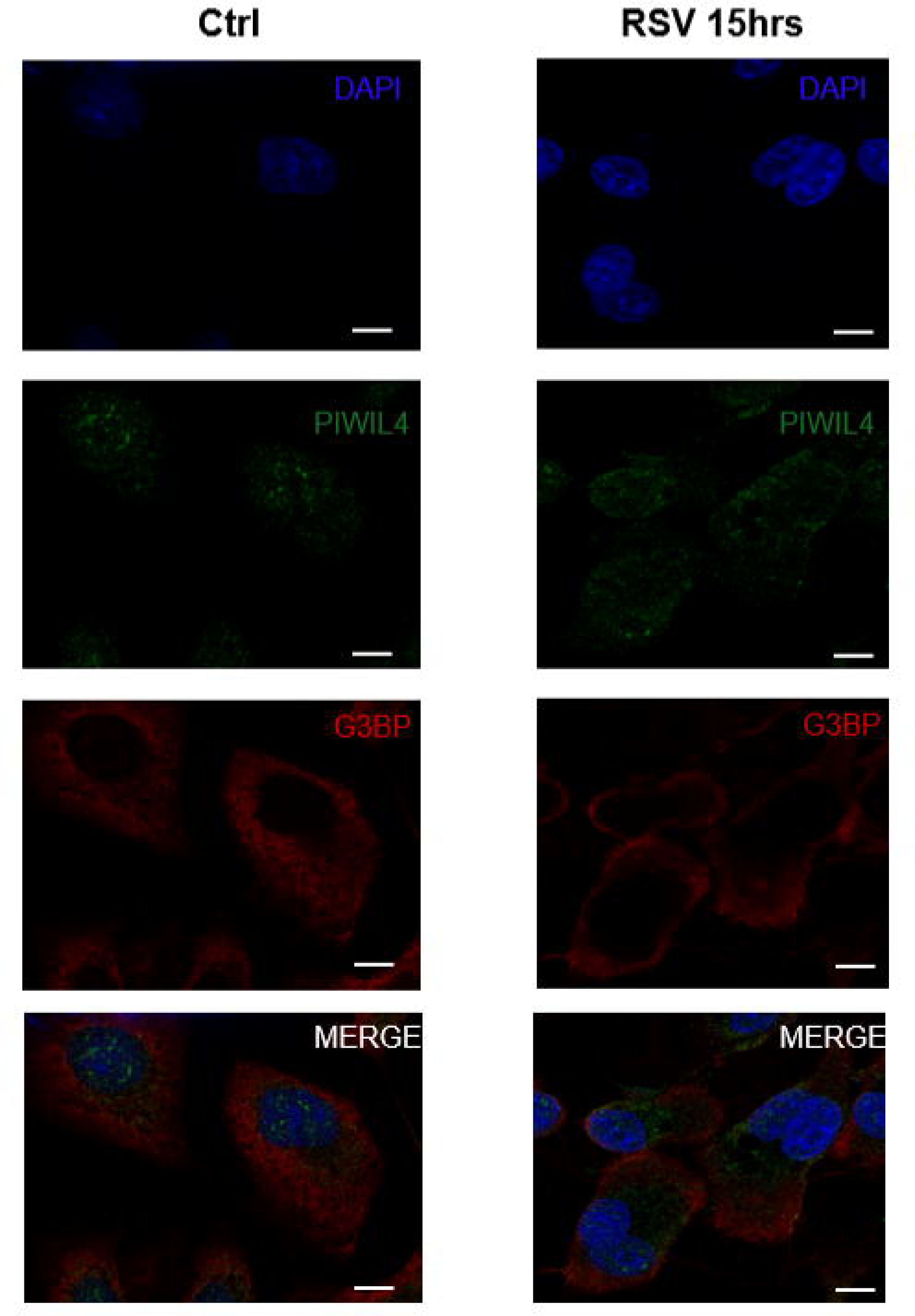
Immunofluorescence detection of PIWIL4 and stress granule marker G3BP in iSAE cells following RSV infection. iSAE cells uninfected (Ctrl) or infected with RSV for 15 hours were stained for PIWIL4 (green) and G3BP (red). Nuclei were counterstained with DAPI (blue), and merged images are shown. Scale bars: 10 µm.

### 2.2 Effect of PIWIL4 knockdown on RSV replication

To determine whether PIWIL4 played a role in regulating RSV replication, we performed siRNA-mediated knockdown (KD) of PIWIL4 in iSAE cells. Cells were transfected with either PIWIL4-targeting or a non-targeting control siRNA, infected with RSV for 15 and 24 ahours and harvested to collect supernatants and extract total RNA. Knockdown efficiency was confirmed by RT-qPCR, which showed significant reduction in PIWIL4 mRNA expression in PIWIL4 siRNA-treated cells compared to non-targeting controls both in basal conditions and following RSV infection, with complete inhibition of the inducible expression observed in response to the infection (**Figure 4A**). To assess viral replication, we performed RSV N gene expression analysis by RT-qPCR in targeting and non-targeting siRNA treated cells infected for 15 hours. PIWIL4 silencing did not significantly reduce cellular RSV N gene levels (**Figure 4B**). A similar experiment was performed to measure RSV titers in the cell culture medium at 24 hours p.i., quantified by plaque assay. Our results showed again no significant difference in RSV replication in PIWIL4 KD cells when compared to non-targeting siRNA-treated cells (**Figure 4C**).

**Figure 4.**
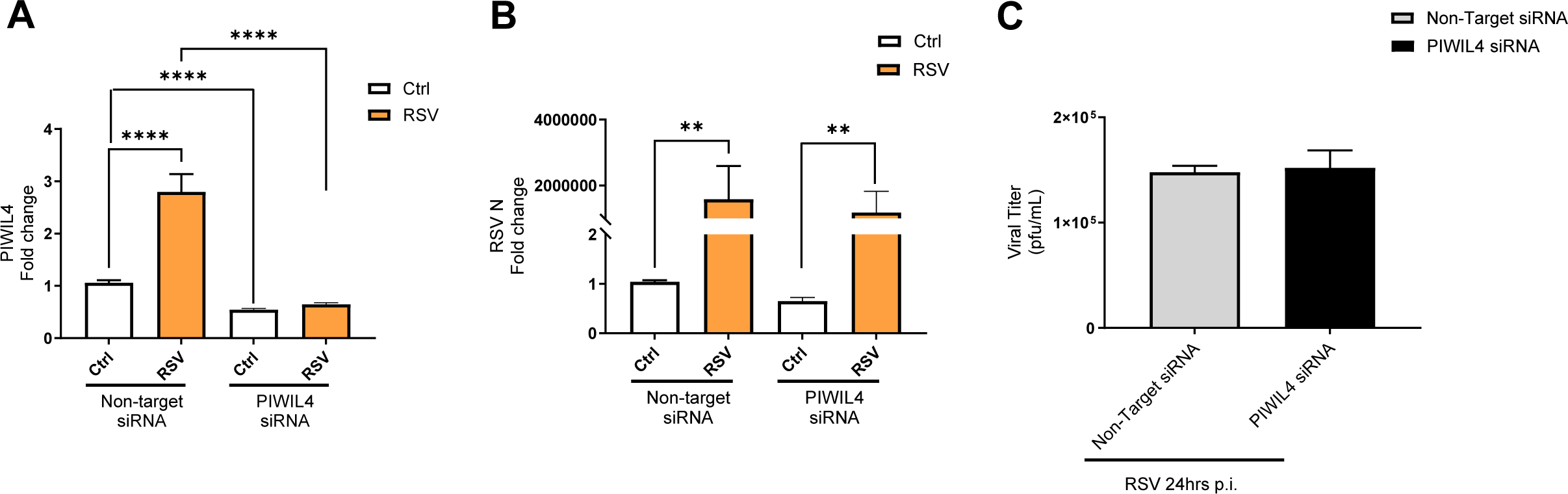
Effect of PIWIL4 silencing on RSV replication in iSAE cells. iSAE cells were transfected with either non-targeting or PIWIL4-targeting siRNA and total RNA was extracted from uninfected cells (Ctrl) or infected with RSV for 15 hours. PIWIL4 mRNA levels **(A)** and RSV N gene expression levels **(B)** were measured by RT-qPCR. Data are means ± SEM (n=2 from two independent experiments run in triplicate; **p<0.01 and ****p<0.0001 determined by one-way ANOVA followed by Tukey’s multiple comparisons test. **(C)** iSAE cells were transfected with either a non-targeting or PIWIL4-targeting siRNA and cell supernatants were harvested from uninfected cells (Ctrl) or infected with RSV for 24 hours to determine viral titers by plaque assay (n=1, from one independent experiment run in triplicate).

### 2.3 Effect of PIWIL4 knockdown on host gene expression

To investigate the effect of PIWIL4 loss on gene expression, we performed transcriptomic profiling of iSAE cells transfected with either PIWIL4-targeting (Targ) or non-targeting (NTarg) siRNA during basal condition (Ctrl) and following RSV infection for 15 hours. Differential gene expression (DEG) and Ingenuity Pathway analysis were conducted to identify genes and the related key pathways affected by PIWIL4 knockdown in both conditions. Using the human Gene Expression Microarrays targeting 27,958 Entrez Gene RNAs, we observed that lack of PIWIL4 resulted in a total of 553 and 252 DEGs significantly upregulated, and 648 and 378 DEGs significantly downregulated in uninfected (Ctrl) conditions and in response to RSV infection, respectively. (**Figure 5A**). The complete list of DEGs in PIWIL-4 deficient cells is included in the data supplements as **Excel file S1**.

**Figure 5.**
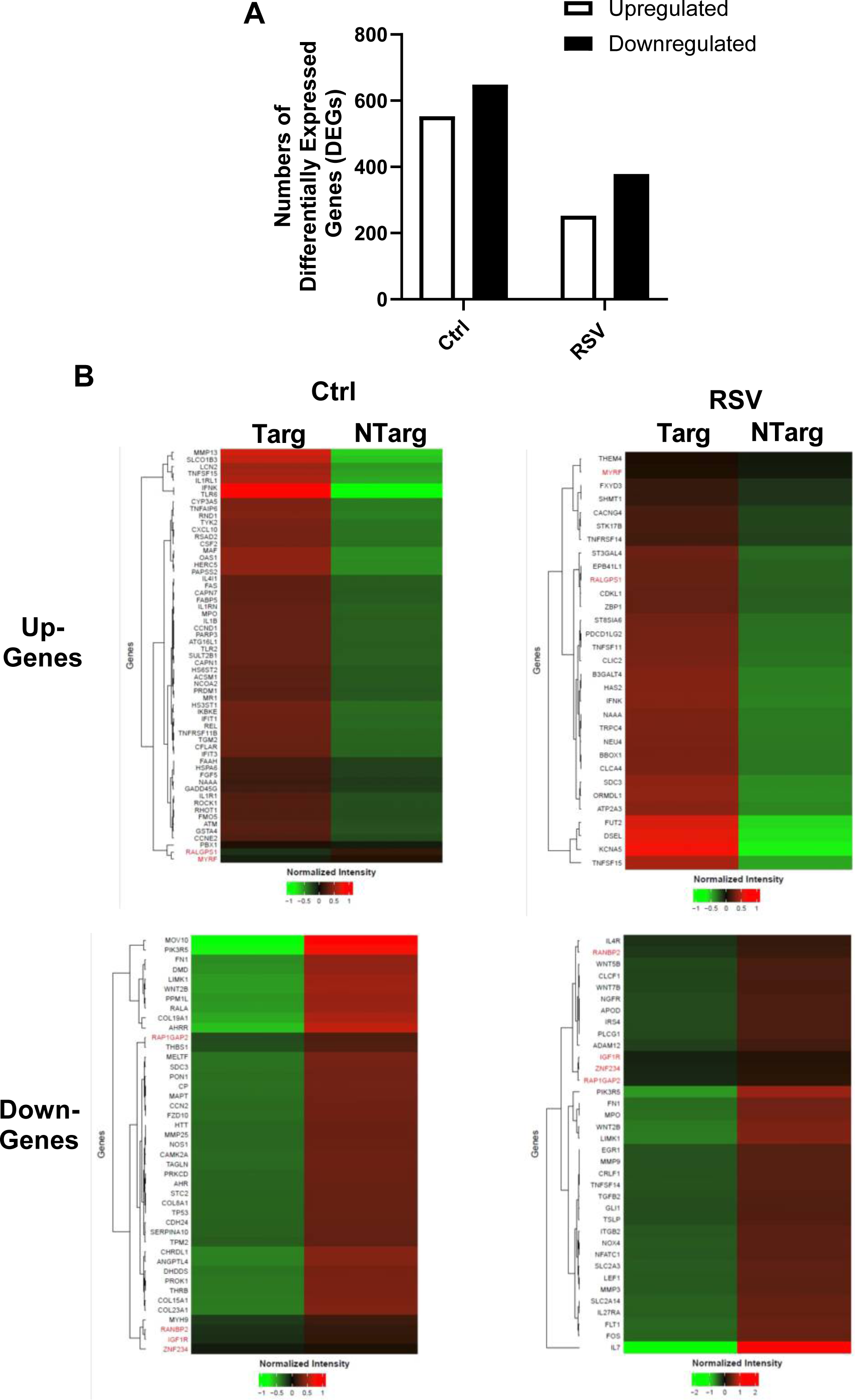
Effect of PIWIL4 silencing on gene expression in iSAE cells. **(A)** iSAE cells were transfected with either non-targeting or PIWIL4-targeting siRNA and total RNA was extracted from cells uninfected (Ctrl) or infected with RSV for 15 hours. Samples from one experiment done in triplicate were pooled and the number of differentially expressed genes (DEGs) were identified by gene array analysis. Number of upregulated (open bars) and downregulated (black bars) DEG in PIWIL4-targeting (Targ) versus non-targeting (NTarg) iSAE cells under Ctrl and RSV infection conditions. Differential expressions were determined using a fold-change cutoff of FC ≥ 1.5. **(B)** Heatmaps showing the expression profiles of significantly differentially expressed genes from the selected IPA-enriched canonical pathways in Tables 1 and 2. Left panels: DEGs under Ctrl conditions; right panels: DEGs under RSV infection. Upper panels: upregulated genes; lower panels: downregulated genes. Red indicates higher relative expression and green indicates lower relative expression (mean-centered log fold-change values).

**Table 1.** Relevant Ingenuity Canonical Pathways of upregulated genes in RSV-infected iSAE PIWIL4 Targeting versus Non-targeting cells.

| **Ingenuity Canonical Pathways**<br>**of upregulated Genes in response to RSV** | **Molecules** |
| --- | --- |
| Carnitine synthesis<br>-log(p-value) = 3.19<br>Ratio=0.5 | *BBOX1, SHMT1* |
| TNFs bind their physiological receptors<br>-log(p-value) = 2.48<br>Ratio=0.1 | *TNFRSF14, TNFSF11, TNFSF15* |
| Sialic acid metabolism<br>log(p-value) = 2.31<br>Ratio=0.09 | *NEU4, ST3GAL4, ST8SIA6* |
| Co-stimulation by the CD28 family<br>log(p-value) = 2.01<br>Ratio=0.04 | *CDKL1, PDCD1LG2, STK17B, THEM4, TNFRSF14* |
| Cardiac conduction<br>log(p-value) = 1.93<br>Ratio=0.03 | *ATP2A3, CACNG4, CLIC2, FXYD3, KCNA5* |
| Glycine biosynthesis I<br>log(p-value) = 1.68<br>Ratio=0.5 | *SHMT1* |
| Sphingolipid metabolism<br>log(p-value) = 1.59<br>Ratio=0.03 | *B3GALT4, FUT2, NEU4, ORMDL1* |
| L-carnitine biosynthesis<br>log(p-value) = 1.51<br>Ratio=0.33 | *BBOX1* |
| Anandamide degradation<br>log(p-value) = 1.51<br>Ratio=0.33 | *NAAA* |
| Glutamate binding, activation of AMPA<br>receptors and synaptic plasticity<br>log(p-value) = 1.38<br>Ratio=0.06 | *CACNG4, EPB41L1* |
| Ion channel transport<br>log(p-value) = 1.37<br>Ratio=0.02 | *ATP2A3, CLCA4, CLIC2, FXYD3, TRPC4* |
| Glycosaminoglycan metabolism<br>log(p-value) = 1.35<br>Ratio=0.03 | *DSEL, HAS2, SDC3, ST3GAL4* |
| CGAS-STING signaling pathway<br>log(p-value) = 1.32<br>Ratio=0.03 | *IFNK, TNFSF11, TNFSF15, ZBP1* |

**Table 2.** Relevant Ingenuity Canonical Pathways of downregulated genes in RSV-infected iSAE PIWIL4 Targeting versus Non-targeting cells.

| **Ingenuity Canonical Pathways**<br>**of downregulated Genes in response to RSV** | **Molecules** |
| --- | --- |
| Pulmonary Fibrosis Idiopathic Signaling Pathway<br>-log(p-value) = 3.04<br>Ratio=0.04 | *EGR1, FN1, FOS, GLI1, LEF1, MMP3, MMP9, NOX4, PIK3R5, TGFB2, WNT2B, WNT5B, WNT7B* |
| Airway Pathology in Chronic Obstructive Pulmonary Disease<br>-log(p-value) = 2.91<br>Ratio=0.06 | *APOD, CLCF1, MMP9, MPO, TGFB2, TNFSF14, TSLP* |
| Pulmonary Healing Signaling Pathway<br>log(p-value) = 2.6<br>Ratio=0.04 | *LEF1, MMP3, MMP9, NFATC1, NGFR, TGFB2, WNT2B, WNT5B, WNT7B* |
| Role of JAK1 and JAK3 in γc Cytokine Signaling<br>log(p-value) = 2.51<br>Ratio=0.07 | *IL4R, IL7, IRS4, PIK3R5, TSLP* |
| HIF1α Signaling<br>log(p-value) = 2.48<br>Ratio=0.04 | *FLT1, MMP3, MMP9, NOX4, PIK3R5, PLCG1, SLC2A14, SLC2A3, TGFB2* |
| Interleukin-4 and Interleukin-13 signaling<br>log(p-value) = 2.25<br>Ratio=0.05 | *FN1, FOS, IL4R, ITGB2, MMP3, MMP9* |
| Regulation of the Epithelial Mesenchymal Transition in Development Pathway<br>log(p-value) = 2.09<br>Ratio=0.05 | *GLI1, LEF1, WNT2B, WNT5B, WNT7B* |
| Airway Inflammation in Asthma<br>log(p-value) = 1.98<br>Ratio=0.09 | *MMP9, TGFB2, TSLP* |
| IL-17 Signaling<br>log(p-value) = 1.94<br>Ratio=0.04 | *CLCF1, FOS, MMP3, MMP9, PIK3R5, TGFB2, TNFSF14* |
| IL-8 Signaling<br>log(p-value) = 1.92<br>Ratio=0.03 | *FLT1, FOS, ITGB2, LIMK1, MMP9, MPO, NOX4, PIK3R5* |
| Inhibition of Matrix Metalloproteases<br>log(p-value) = 1.74<br>Ratio=0.07 | *ADAM12, MMP3, MMP9* |
| Role of JAK family kinases in IL-6-type Cytokine Signaling<br>log(p-value) = 1.57<br>Ratio=0.05 | *CLCF1, FOS, IL27RA, TGFB2* |
| Interleukin-12 family signaling<br>log(p-value) = 1.46<br>Ratio=0.1 | *CRLF1, IL27RA* |

To further elucidate the functional impact of PIWIL4 knockdown on the host transcriptomics, we performed Ingenuity Pathway analysis of up- and downregulated DEGs identified in uninfected and infected PIWIL4 knockdown cells. DEGs associated with the most relevant pathways are shown in the heatmaps in **Figure 5B**. Pathways enriched among the upregulated genes in response to RSV infection were predominantly related to innate immunity, stress response and metabolic pathways, as listed in **Table 1** (genes included in **Figure 5B, upper right panel**). Immune-related pathways included genes associated with TNF binding receptors (TNFRSF14, TNFSF11, TNFSF15), cGAS-STING signaling pathway (IFNK, TNFSF11, TNFSF15, ZBP1) and the CD28 T-cell co-stimulatory pathway (CDKL1, PDCD1LG2, STK17B, THEM4, TNFRSF14), while biosynthetic pathways included genes related to sialic acid, sphingolipid and glycosaminoglycan metabolism that have been shown to play a role in viral-induced intracellular signaling, virus cell attachment and other steps of viral replication^25–27^. Conversely, pathways enriched among the downregulated genes in PIWIL4-deficient infected cells were predominantly associated with cytokine signaling, including IL-4, IL-6, IL-8, IL-12, IL-13 and IL-17, Jak/STAT signaling, as well as airway inflammation and remodeling, as listed in **Table 2** (genes included in **Figure 5B, lower right panel**). In uninfected (Ctrl) condition, Aryl Hydrocarbon Receptor (AhR), Nuclear Factor-kappa B inhibitor epsilon (IKBε), HMGB1, apoptosis and pyroptosis signaling were the key canonical pathways significantly associated with the upregulated genes (genes included in **Figure 5B, upper left panel**). IPA of the downregulated genes included networks related to mitochondrial dynamics, protein modification, extracellular matrix remodeling, and calcium-mediated signaling (genes included in **Figure 5B, lower left panel**). The full list of the enriched pathways of up- and downregulated DEGs in uninfected (Ctrl) and RSV infected conditions is included in the data supplement as **Excel files S2**.

To further evaluate the impact of PIWIL4 knockdown on the RSV-induced epithelial innate response, we measured the levels of key pro-inflammatory cytokines, chemokines and growth factors in culture supernatants of uninfected and infected cells. Using a human multi-plex array, we found that PIWIL4 knockdown led to a moderate increase of IL-6 and a concomitant reduction of IL-8, G-CSF, and Vascular Endothelial Growth Factor (VEGF) secretion in response to RSV infection (**Figure 6**).

**Figure 6.**
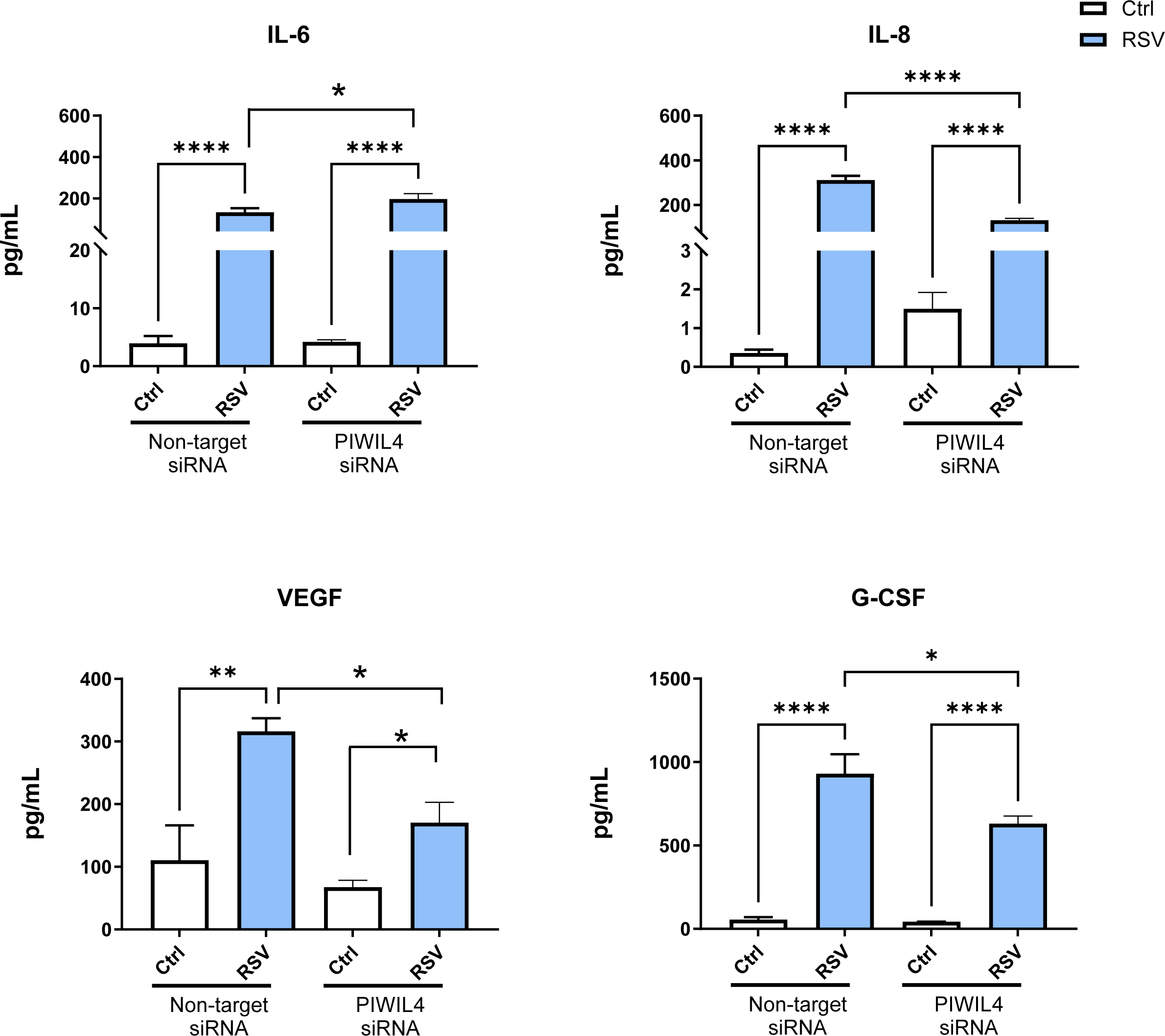
Effect of PIWIL4 silencing on immune mediator production in RSV-infected iSAE cells. Levels of immune mediators were measured by 48-human multi-Plex array in supernatants from cells transfected with non-targeting siRNA or PIWIL4-Targeting siRNA and infected with RSV for 24 hours. Data are means ± SEM; (n=2, from two independent experiments run in triplicate; *p<0.05, **p<0.01, ****p<0.0001 by one-way ANOVA followed by Tukey’s multiple comparisons test).

### 2.4 Effect of PIWIL4 knockdown on piRNA expression in control and RSV-infected airway epithelial cells

To explore the role of PIWIL4 loss on piRNA expression in airway epithelial cells, we performed piRNA profiling of iSAE cells transfected with either PIWIL4-targeting (Targ) or non-targeting (NTarg) siRNA during basal condition (Ctrl) and following RSV infection for 15 hours, using the Arraystar Human piRNA Microarray (HG19) that targets 23,677 unique piRNAs. As illustrated in **Figure 7A**, there were 9,948 piRNAs identified in both uninfected and infected NTarg samples, with 975 piRNAs detected only in RSV-infected cells and 247 only in uninfected (Ctrl). In Targ samples, 650 piRNAs were present only in RSV, 566 only in controls, and 9,908 were common to both groups.

**Figure 7:**
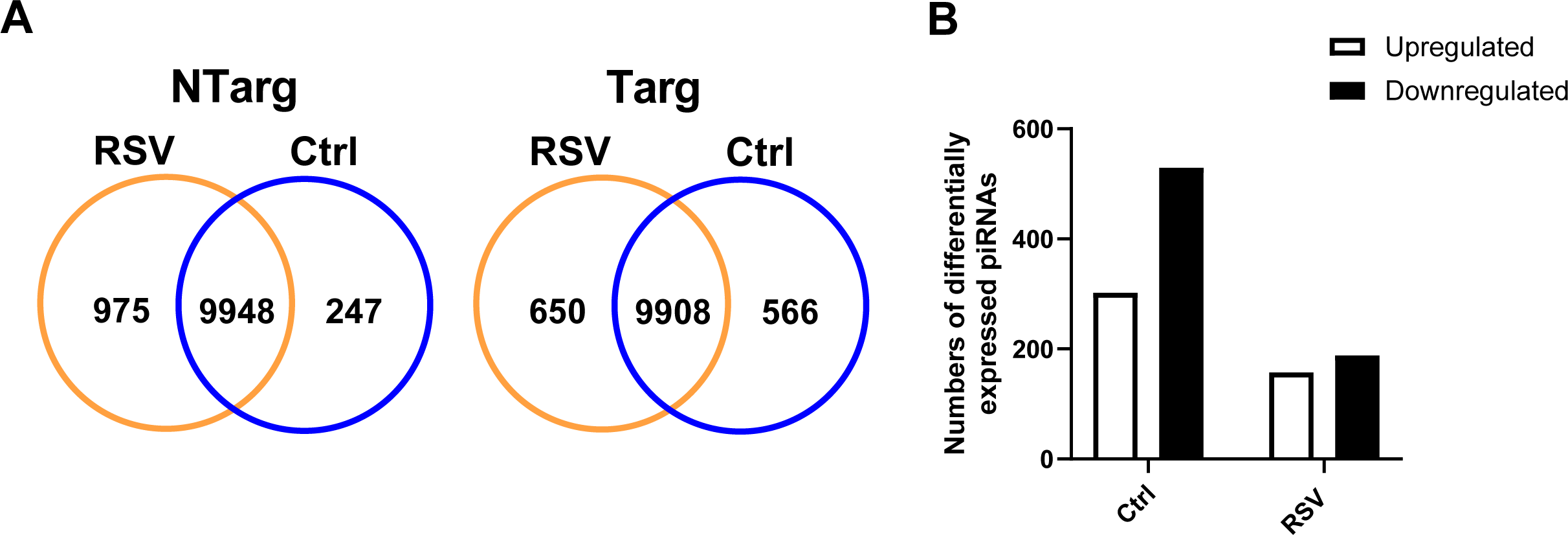
Detection and differential expression of piRNAs in PIWIL4-silenced iSAE cells following RSV infection. iSAE cells were transfected with either non-targeting or PIWIL4-targeting siRNA and total RNA was extracted from uninfected cells (Ctrl) or infected with RSV for 15 hours. Samples from one experiment done in triplicate were pooled and the number of differentially expressed genes (DEGs) were identified by piRNA array analysis. **(A)** Venn diagrams illustrating the distribution of piRNAs detected among the four experimental groups: non-targeting control (NTarg Ctrl), non-targeting RSV-infected (NTarg RSV), PIWIL4 targeting control (Targ Ctrl), and PIWIL4 targeting RSV-infected (Targ RSV) groups. Numbers within each section indicate piRNAs detected exclusively in one condition or shared between conditions. For NTarg cells, 975 piRNAs were detected only in RSV-infected samples, 247 only in controls, and 9,948 for both conditions. For Targ cells, 650 piRNAs were unique to RSV-infected samples, 566 unique to controls, and 9,908 were detected in both. **(B)** Bar graphs display the numbers of differentially upregulated (open bars) or downregulated (black bars) expressed piRNAs comparing PIWIL4 targeting versus non-targeting conditions within either the control or RSV-infected condition. Differential expressions were determined using a fold-change cutoff of FC ≥ 1.5.

After quantile normalization and subsequent data processing, a total of 302 piRNAs were identified as differentially upregulated in iSAE under basal conditions and 157 piRNAs upregulated in response to RSV infection, when comparing iSAE PIWIL4 targeting versus non-targeting groups (**7B, open bars**). Notably, a larger number of piRNAs were significantly downregulated upon PIWIL4 knockdown, with 529 piRNAs differentially expressed in control condition and 188 following RSV infection (**7B, black bars**). The complete list of piRNAs whose expression differs in PIWIL-4 deficient cells is included in the data supplements as **Excel file S3**. Altogether, this data indicates that PIWIL4 plays a critical role in maintaining the expression of a substantial portion of the piRNA repertoire in airway epithelial cells, which is further modulated during viral challenge.

To validate the expression patterns identified in the piRNA array analysis, we performed RT-qPCR for ten differentially expressed piRNAs selected among the top twenty either upregulated or downregulated in iSAE cells control or infected with RSV for 15 hours. Among the upregulated piRNAs in virus-infected KD cells compared to non-targeting controls, piR-37912 (DQ599846) showed the highest induction, with a greater than threefold increase (**Figure 8A**, red-shaded areas). Four additional piRNAs, piR-58759 (DQ591647), piR-37822 (DQ599756), piR-43080 (DQ574968) and piR-30426 (DQ570314) were upregulated by more than two-fold, three piRNAs, piR-39412 (DQ601346), piR-30908 (DQ570796), piR-36946 (DQ598880) exhibited more than one and half fold change and two piRNAs, piR-39690 (DQ601624) and piR-52554 (DQ585442), were upregulated by more than one-fold in PIWIL4-targeting (Targ) compared to non-targeting control (NTarg) virus-infected iSAE cells. Among the ten upregulated piRNAs in control condition (**Figure 8A**, blue-shaded areas), five piRNAs, piR-60909 (DQ594797), piR-43076 (DQ574964), piR-61909 (DQ595797), piR-40620 (DQ572508), and piR-36816 (DQ598750), exhibited the highest induction, with fold changes of 3.63, 2.97, 2.30, 2.11, and 2.46, respectively. The remaining five piRNAs, piR-60168 (DQ594056), piR-58815 (DQ591703), piR-59558 (DQ592446), piR-56637 (DQ589525), and piR-60902 (DQ594790) were also upregulated, showing more moderate increases ranging from 1.38 to 1.94-fold.

**Figure 8.**
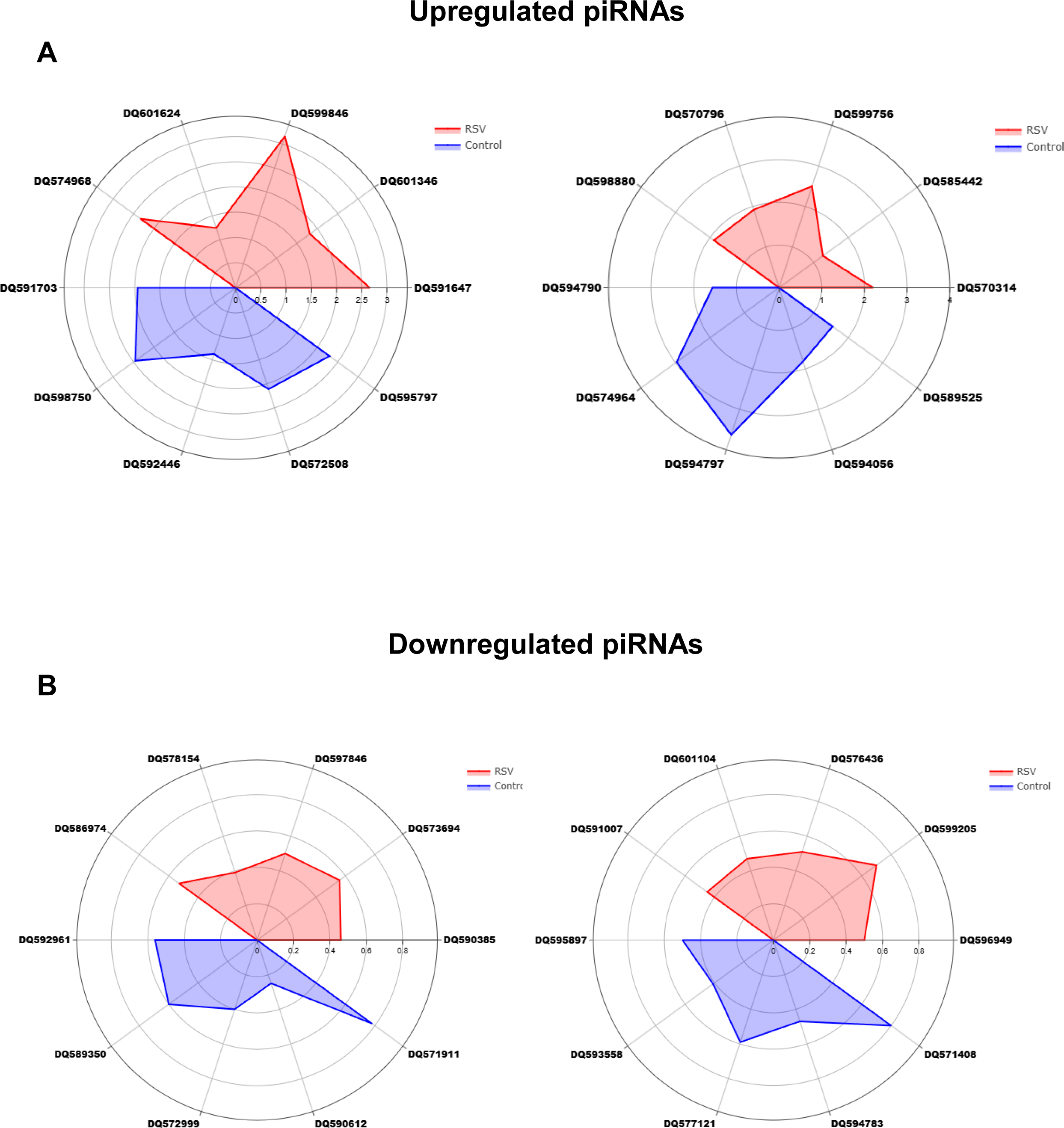
Validation of selected upregulated (A) and downregulated (B) piRNAs in PIWIL4-targeting iSAE cells following RSV infection. Radar plots depicting the expression profiles of twenty selected piRNAs in PIWIL4-targeting (Targ) versus non-targeting (NTarg) iSAE cells either infected with RSV for 15 hours (red shaded area) or uninfected (control) (blue shaded area). piRNA expression levels were measured by RT-qPCR and calculated as fold changes relative to NTarg control cells using the 2^−ΔΔCT method. Data are normalized to U6 small nuclear RNA and presented as mean ± SEM (n=2, from two independent experiments run in triplicate).

Among the ten selected differentially downregulated piRNAs in RSV-infected PIWIL4-KD cells (**Figure 8B**, red-shaded area), piR-46266 (DQ578154), piR-58119 (DQ591007), piR-57497 (DQ590385), piR-39170 (DQ601104), and piR-44548 (DQ576436) displayed marked reductions, with fold changes of 0.39, 0.45, 0.46, 0.47, and 0.51, respectively. Additional downregulated piRNAs included piR-35015 (DQ596949), piR-54086 (DQ586974), piR-41806 (DQ573694), piR-37271 (DQ599205), and piR-35912 (DQ597846), showing fold changes between 0.50 and 0.70. Among the ten selected differentially downregulated piRNAs in basal conditions, piR-57724 (DQ590612), piR-41111 (DQ572999), piR-33670 (DQ593558), and piR-60895 (DQ594783) exhibited the strongest reductions, with fold changes of 0.25, 0.40, 0.41, and 0.47, respectively. The remaining downregulated piRNAs, piR-33073 (DQ592961), piR-62009 (DQ595897), piR-56462 (DQ589350), piR-45233 (DQ577121), piR-32023 (DQ571911), and piR-31520 (DQ571408) ranged in fold change from 0.50 to 0.80 (**Figure 8B**, blue-shaded area).

Next, we characterized the predicted targets of piRNAs differentially expressed in RSV-infected and control iSAE cells knockdown for PIWIL4 and integrated them with transcriptomic changes observed in PIWIL4-silenced cells found by gene arrays. The complete list of predicted targets and targets found in the gene arrays for the upregulated and downregulated piRNAs in control and infected PIWIL4-silenced cells is included in the data supplements as **Excel file S4 and S5**, respectively. Most of the differentially expressed piRNAs had multiple predicted gene targets, however, we found minimal overlap between those, and the identified genes experimentally altered in the same conditions. Among the differentially upregulated piRNAs in response to RSV infection, we found twelve piRNAs with genes common to both transcriptomic data and predicted targets (**Table 3**). These genes were: Dynamin 1 Pseudogene 47 (DNM1P47), SPATA31 subfamily C member 2 (SPATA31C2), Galactosidase Beta 1 Like (GLB1L), ST6 N-acetylgalactosaminide alpha-2,6-sialyltransferase 5 (ST6GALNAC5), scavenger receptor family member expressed on T cells 1 (SCART1), sphingosine-1-phosphate phosphatase 2 (SGPP2), TBC1 domain family member 5 (TBC1D5), obscurin (OBSCN), NIPA like domain containing 2 (NIPAL2), Zinc Finger Protein 652 (ZNF652), Homeobox Protein CP19 (HOXC4), Acetyl-CoA Carboxylase Alpha (ACACA), Cyclin M2 (CNNM2), TRAF2 and NCK Interacting Kinase (TNIK), and Neurotrimin (NTM). Among the differentially downregulated piRNAs in response to RSV infection, we found fifteen piRNAs with genes common to both transcriptomic data and predicted targets (**Table 4**). The overlapping genes were: Ral GEF with PH domain and SH3 binding motif 1 (RALGPS1), RAS and EF-hand domain containing (RASEF), Golgin A8 family member M (GOLGA8M), NME/NM23 family member 7 (NME7), phospholipase C gamma 1 (PLCG1), immunity related GTPase Q (IRGQ), Dynamin 1 pseudogene 47 (DNM1P47), RAN binding protein 2 (RANBP2), endoplasmic reticulum protein 44 (ERP44), Nischarin (NISCH), egl-9 family hypoxia inducible factor 3 (EGLN3), Fc gamma receptor IIb (FCGR2B), Rho GTPase-activating protein 28 (ARHGAP28), suppressor of cancer cell invasion (SCAI), phosphodiesterase 4D (PDE4D), sperm antigen with calponin homology and coiled-coil domains 1 like (SPECC1L), RNA polymerase II subunit J4 (POLR2J4) and methylthioadenosine phosphorylase (MTAP).

**Table 3.**
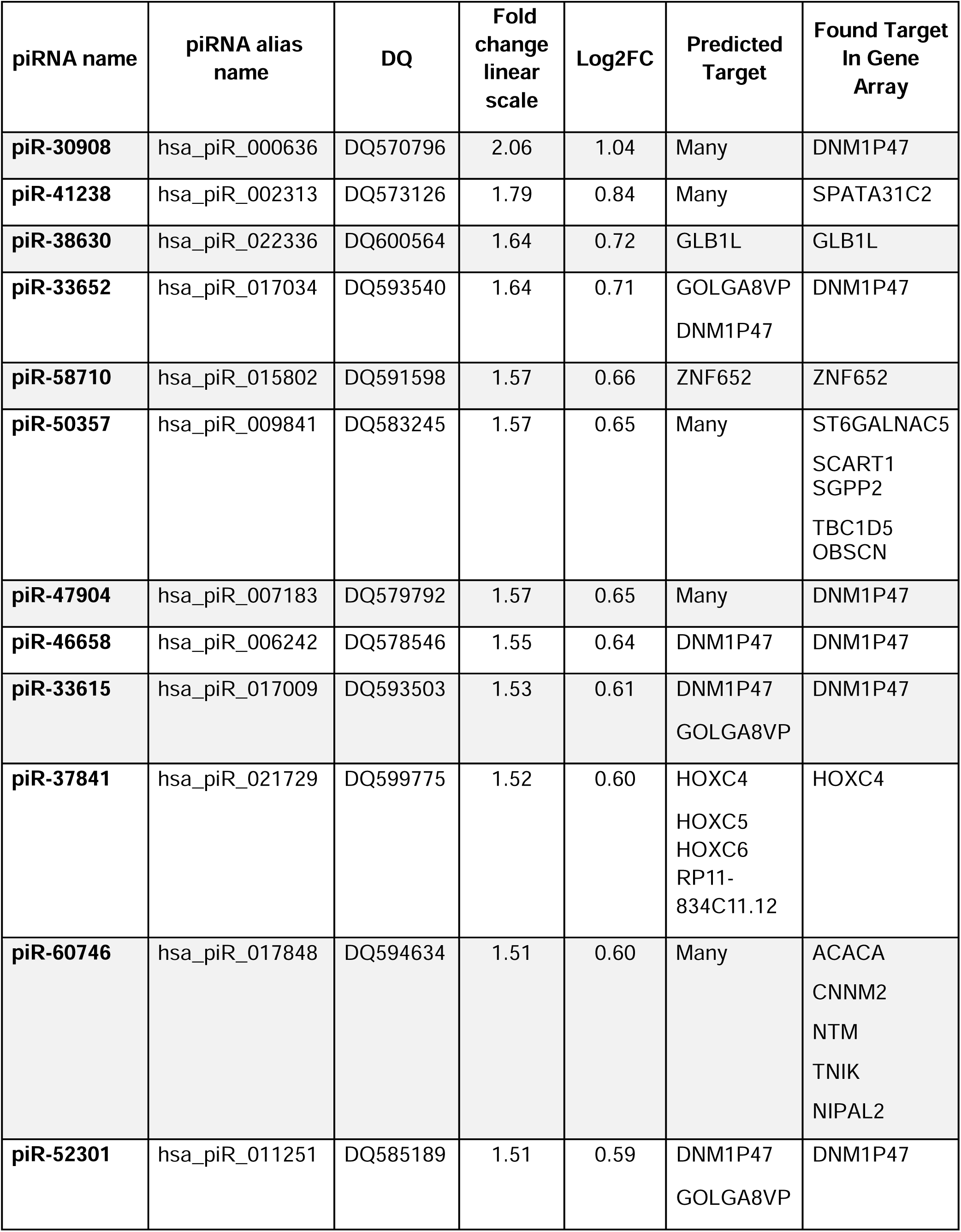
List of all differentially upregulated piRNAs (≥1.5-fold) with an identified target in the gene array in RSV-infected iSAE cells, comparing PIWIL4-targeting versus non-targeting conditions with an identified target (*p ≤ 0.05). Shown are the piRNAs, their predicted targets, and the target identified in the gene array analysis. “DQ” denotes the accession number of each piRNA in the NCBI database.

**Table 4.**
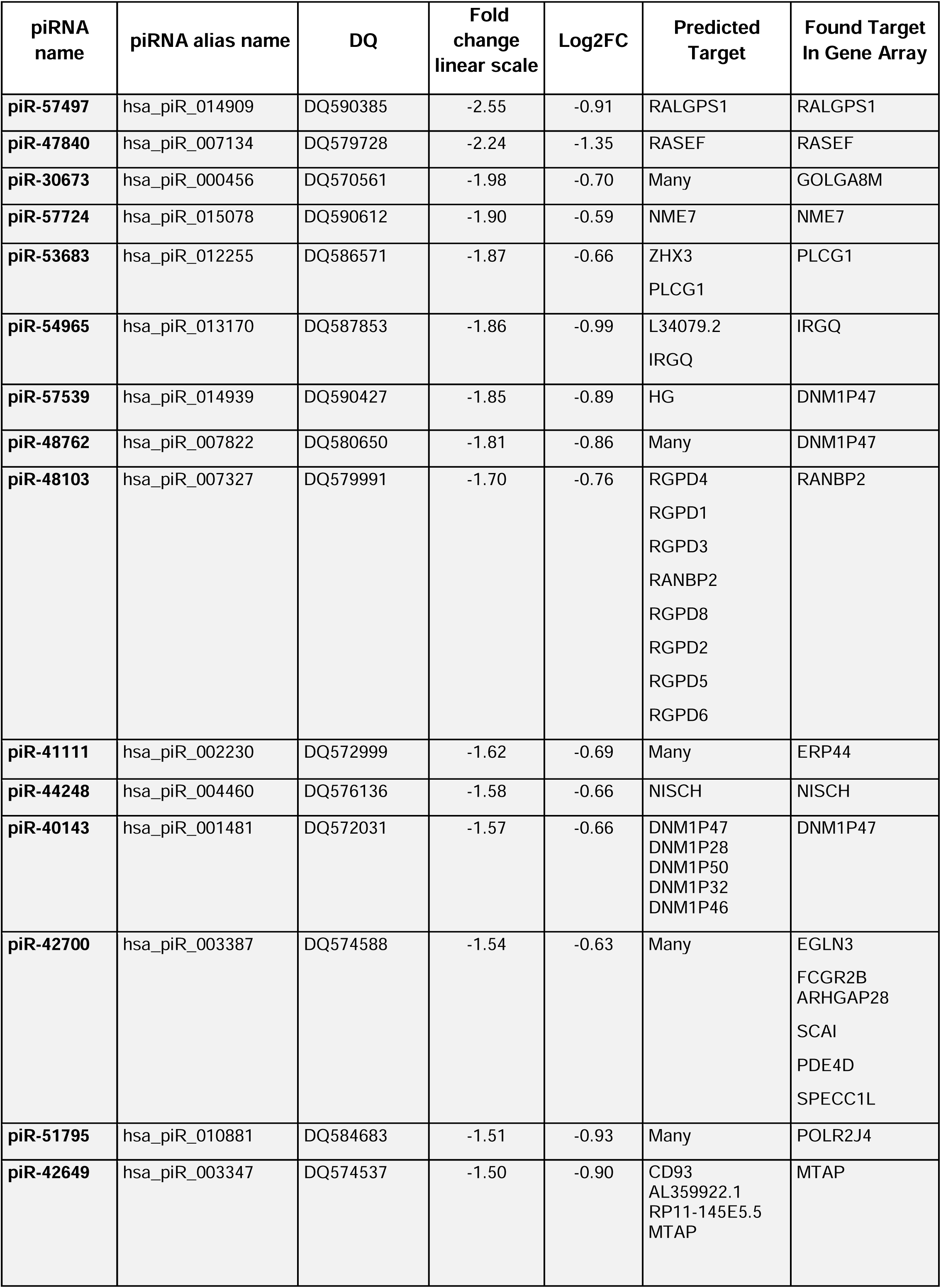
List of all differentially downregulated piRNAs (≥1.5-fold) with an identified target in the gene array in RSV-infected iSAE cells, comparing PIWIL4-targeting versus non-targeting conditions with an identified target (*p ≤ 0.05). Shown are the piRNAs, their predicted targets, and the target identified in the gene array analysis. “DQ” denotes the accession number of each piRNA in the NCBI database.

## 3. Methods

### RSV Preparation

The RSV Long strain was grown in Hep-2 cells and purified by centrifugation on discontinuous sucrose gradients as described elsewhere ^28^. The virus titer of the purified RSV pools was 8–9 log10 plaque forming units (PFU)/mL using a methylcellulose plaque assay. No contaminating LPS or cytokines were found in these sucrose-purified viral preparations ^29^. Virus pools were aliquoted, quick-frozen on dry ice/alcohol and stored at −80 °C until used.

### Cell cultures, RSV infection and polyinosinic-polycytidylic acid (Poly I:C) treatment

Primary human small airway epithelial (SAE) cells, derived from terminal bronchioli of two different cadaveric donors (25- and 38-years old donors), were purchased from Lonza Inc., San Diego, CA. Immortalized SAE (iSAE) cells were established by transducing primary cells with human telomerase and cyclin-dependent kinase-4 retrovirus constructs, as described in ^21^. Both cell types were grown in growth medium, containing 7.5 mg/ml bovine pituitary extract (BPE), 0.5 mg/ml hydrocortisone, 0.5 μg/ml hEGF, 0.5 mg/ml epinephrine, 10 mg/ml transferrin, 5 mg/ml insulin, 0.1 μg/ml retinoic acid, 0.5 μg/ml triiodothyronine, 50 mg/ml gentamicin and 50 mg/ml bovine serum albumin. SAE and iSAE cells were switched to basal media (no supplements added) 4 hours prior to RSV infection or treatment with poly(I:C) (10 μg/ml, Sigma, St. Louis, MO). At 90%–95% confluence, cell monolayers were infected with RSV at multiplicity of infection (MOI) of 3 for all the described experiments. An equivalent amount of 30% sucrose solution was added to uninfected SAE cells as a control (uninfected cells). Cells or supernatants were collected at 6-, 15- and 24-hours RSV post-infection (p.i.) or post-poly I:C treatment, depending on the type of assay and analysis performed.

### siRNA silencing

Three custom-synthesized human siRNA sequences, PIWIL4 siRNA duplex, 21mer, HPLC Purification, (VC30002, Sigma, St. Louis, MO), were used as a mixture to silence gene PIWIL4 expression at a final concentration of 300 nM. Non-Targeting sequences (SIC001-10NMOL, Sigma, St. Louis, MO) were used as negative control at the same concentration. The PIWIL4 sequences of siRNA are shown in the following table.

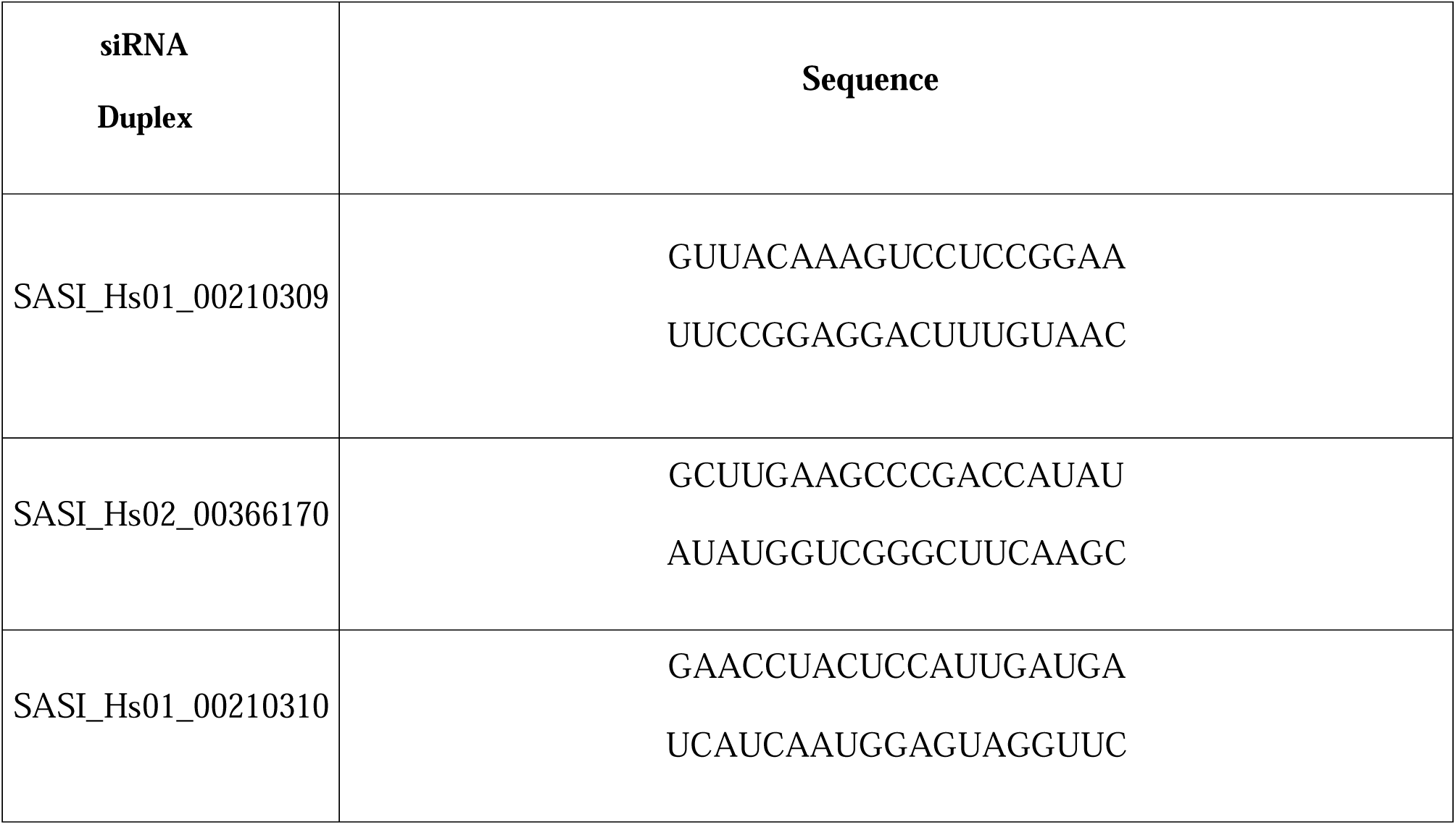

The siRNAs were transfected by electroporation using the Amaxa Biosystem, Gaithersburg, MD, and the Amaxa Basic Nucleofector Kit (Cat # VPI-1005) for iSAE cells, according to manufacturer instructions, to allow high transfection efficiency. At 24-30 hours post-transfection, cells were infected with RSV and harvested at 15 or 24 hours p.i. to extract total RNA and/or collect the supernatants.

### Total RNA extraction and Reverse transcription quantitative PCR (RT-qPCR)

Total RNA was extracted using ToTALLY RNA kit from Ambion (Life Technologies, Carlsbad, CA). RNA concentration was quantified using a Nanodrop Spectrophotometer (Nanodrop Technologies, Wilmington, DE) and integrity was assessed using standard denaturing agarose gel electrophoresis. RNA was used for gene and piRNA arrays (see details below), and RT-qPCR analysis. cDNA was prepared using iScript Reverse Transcription Supermix (Bio-Rad, Hercules, CA) and qPCR was performed with human primer PIWIL4 (Integrated DNA Technologies, Coralville, IA) and SYBR Green master mix (ABclonal, Woburn, MA). Data was analyzed using the 2^-ΔΔCt^ method and 18S rRNA (Integrated DNA Technologies, Coralville, IA) was used as endogenous control. For the viral replication quantification, RSV N gene was amplified using specific primers, as described ^30^. Primer sequences are available upon request.

### Western blot

Whole-cell lysates were prepared using SDS sample buffer (New England BioLabs, Ipswich, MA) as described in ^31^. Equal volumes (30 μL) of proteins were separated by SDS-PAGE and transferred onto polyvinylidene difluoride (PVDF) membrane. Nonspecific binding was blocked by immersing the membrane in Tris-buffered saline-Tween (TBST) blocking solution containing 5% skim milk powder. After blocking, the membranes were incubated with the primary antibody overnight at 4°C, followed by the appropriate secondary antibody diluted in TBST for 1h at room temperature. Proteins were detected using enhanced chemiluminescence. The primary antibody used for PIWIL4 (Abcam, ab87939). β-Actin was used as loading control protein to normalize the target proteins expressions from whole cell extracts (Santa Cruz, sc-47778).

### Confocal Microscopy

iSAE cells (∼2 × 10^5^ cells per well) were seeded on coverslips in 12 well plates in SAE growth medium containing 10% FBS and incubated overnight at 37 °C. The cells were then uninfected or infected with RSV for 15 h. The cells were fixed with 4% paraformaldehyde solution for 20 min at room temperature and then washed twice with PBS. The cells were blocked and permeabilized with 0.1% Triton X-100 and 3% BSA in PBS for 30 min at room temperature and washed. The cells were then incubated with primary antibodies: rabbit anti-PIWIL4, 1:500 dilution, (Abcam, ab180867) and mouse anti-G3BP, 1:600 dilution, (Abnova, H00010146-M01) in 3% BSA in PBS for 1h at room temperature. The cells were washed three times with PBS and incubated with secondary antibodies: anti-rabbit Alexa Flour 488 IgG (H+L), 1:4000 dilution, (Invitrogen, A-11008) and anti-mouse Alexa Flour 594 IgG (H+L), 1:4000 dilution, (Invitrogen, A-11005) overnight at 4 °C. The cells were washed three times with PBS, and the coverslips were mounted onto glass slides using a drop of ProLong Gold Antifade reagent with DAPI (Invitrogen), dried and kept overnight at 4 °C. Imaging was performed on a Zeiss LSM 510 Meta confocal laser scanning microscope at excitation/emission wavelengths of 488/505–550 nm (for green fluorescence), 561/570–620 nm (for red fluorescence), and 405/425–475 nm (for DAPI, colored blue). Pictures of the fluorescence patterns were merged to visualize colocalization using ImageJ software version ImageJ 1.54g downloaded from the ImageJ official website.

### Multiplex cytokine analysis

Cytokines, chemokines and growth factors were measured as pg/mL using the 48-target human multi-Plex panel (Bio-Rad Laboratories, Hercules, CA, USA) according to the manufacturer’s instructions and as previously published ^32, 33^.

### Gene microarray and canonical pathway analysis

Gene expression analysis was performed by Arraystar Inc. (Rockville, MD, USA). The sample preparation and array hybridization were performed based on the manufacturer’s standard protocols. Briefly, total RNA extracted from non-targeting and PIWIL4 siRNA-treated iSAE cells, control or infected with RSV for 15 hours, was transcribed into fluorescent cRNA using the manufacturer’s Agilent’s Quick Amp Labeling protocol (version 5.7, Agilent Technologies). The labeled cRNAs were hybridized onto the Whole Human Genome Oligo Microarray (4 x 44K, Agilent Technologies). After washing, the arrays were scanned by the Agilent Scanner G2505C. Agilent Feature Extraction software (version 11.0.1.1) was used to analyze acquired array images. Quantile normalization and subsequent data processing were performed using the GeneSpring GX v12.1 software (Agilent Technologies). After quantile normalization of the raw data, genes that had flags in Detected (“All Targets Value”) for at least 1 out of 4 samples were chosen for further data analysis. Differentially expressed genes were identified through Volcano filtering. Ingenuity Pathway Analysis was used to identify relevant biological pathways altered by PIWIL4 knockdown. p values were calculated by Fisher’s exact test and enrichment scores are log10(P value). Excel files of original data are provided as supplemental material. Canonical pathway analysis was performed using *QIAGEN Ingenuity Pathway Analysis (IPA)*. The methods and algorithms implemented in IPA have been previously described^34^. Genes with an absolute fold change ≥ 1.5 and a p-value < 0.05 were included. Pathways significantly associated with the dataset were identified from the IPA canonical pathway library. Statistical significance was assessed using a right-tailed Fisher’s exact test, with pathways considered significant at p ≤ 0.05 (i.e., –log (p-value) ≥ 1.3). IPA also provided: a ratio, defined as the number of dataset molecules mapping to the pathway divided by the total molecules in that pathway; and a z-score, used to predict pathway activation or inhibition based on the expression patterns relative to the IPA Knowledge Base. Heatmaps were generated in R 4.4.3 using normalized, mean-centered gene expression data with hierarchical clustering applied to genes.

### piRNA array and piRNA RT-qPCR

piRNA array analysis was performed by Arraystar Inc. (Rockville, MD, USA) using the Arraystar Human piRNA Microarray (HG19), which targets 23,677 unique piRNAs and data analyzed as in ^13^. Briefly, 600 ng of total RNA, coming from the same samples used for the gene arrays described above, was reversed transcribed into cDNA in 1X RT buffer, 50 nM RT primers, 250 nM dNTPs, 0.6 U RNase Inhibitor, and 3 U M-MuLV Reverse Transcriptase in a 20 μl reaction volume, as previously published ^13^. Analysis of the differentially expressed piRNA was performed using the PDL:Stats and Statistics:PointEstimation modules from CPAN (<u>PDL-Stats</u>; <u>PDL - the Perl Data Language -</u> metacpan.org). Fold change of the differentially expressed piRNAs were calculated as linear average or Log2 average ratio. To validate piRNA results, real-time qPCR was performed by Arraystar Inc., data was analyzed using the 2^-ΔΔCt^ method and the relative amount of the target piRNA was normalized using U6 as internal control. piRNA primer sequences are available upon request.

### Predicted targets of piRNAs

Prediction of potential piRNA targets was performed by combining piRNA alignments with lists of genomic features, using the bedtools package. To annotate the potential targets of the identified piRNAs, we used the genomic coordinates of the piRNAs, as available from the piRNAQuest resource, http://dibresources.jcbose.ac.in/zhumur/pirnaquest2/^35^. We used the UCSC liftOver tool ^36^ and BEDTools^37^ to identify overlaps of piRNA sequences with genomic features, as in the piRNAdb database, considering overlaps on both strands of the genomic DNA. We verified the implementation by comparing selected results against piRNAdb.

### Statistical analysis

A one-way ANOVA followed by Tukey’s multiple comparisons test was performed using GraphPad Prism v4 (GraphPad Software) for specific experimental data analysis. Significance is indicated as a p value of <0.05 (*) or 0.01 (**) or 0.001 (***) or 0.0001 (***) and data are presented as mean ± SEM. using GraphPad Prism v4 (GraphPad Software).

## 4 Discussion

RSV is a leading cause of hospitalization and death in children and in older adults, resulting in 33.1 million cases and 3.2 million hospitalizations in children < 5 years old ^38^, and with an estimated 177,000 hospitalizations and 14,000 deaths among the elderly in the United States^39^. PIWI proteins are a specialized subfamily of the Argonaute protein (AGO), best known for their role in germline development and fertility, RNA silencing and gene regulation ^4, 15^. In humans, four PIWI-clade proteins have been identified: PIWIL1 (HIWI), PIWIL2 (HILI), PIWIL3, and PIWIL4 (HIWI2) ^6^. Their interacting partners, PIWI-interacting RNAs (piRNAs), represent the largest class of small non-coding RNAs expressed in animal cells and can originate from diverse sources, including transposons, mRNAs and long non-coding RNAs (lncRNAs), small nucleolar RNAs (snoRNAs), and tRNA-derived fragments (tRFs). piRNAs, together with PIWI proteins, play a critical role in maintaining genome integrity by suppressing insertional mutations caused by transposable elements ^4^. Although originally characterized in germline cells, growing evidence indicates that PIWI proteins and piRNAs also regulate gene expression in somatic cells ^7, 15^. The biogenesis and function of piRNAs involve PIWI proteins, Argonaute-3 (AGO3), and Aubergine (AUB). PIWI proteins are present in both somatic and germ cells, while AGO3 and AUB are primarily restricted to germ cells. piRNA precursors are exported from the nucleus into the cytoplasm, where they undergo processing into smaller sequences that bind PIWI proteins to form piRNA-PIWI complexes. These complexes have been implicated in silencing of retrotransposons and other genetic elements, particularly during spermatogenesis ^10^. Although the mechanisms of piRNA generation are well defined, to date, the role of PIWI proteins and piRNA pathway in human airway epithelial cell responses to viral infections is still unknown. Our findings demonstrated for the first time that PIWIL4 is significantly upregulated at both the mRNA and protein levels in primary SAE cells following RSV infection and poly I:C stimulation, a proxy of viral infection, suggesting a role for this PIWI protein in the host response to viruses. RT-qPCR and Western blot analyses showed a time-dependent increase in PIWIL4 expression, with a peak around 24 hours after both RSV infection and poly(I:C) treatment. Recently, MIWI2 (the murine homolog of PIWIL4) was found to be expressed in lung epithelial cells of adult mice and was induced in response to bacterial pneumonia ^11^. A previous study in synovial fibroblasts cultured from rheumatoid arthritis patients also showed inducible PIWIL4 in response to TNFα+IL1β/TLR-ligands ^40^. Similarly, PIWIL4 levels increased in a time-dependent manner during trans retinoic acid (RA)-mediated neuronal differentiation between days 15 and 18 post-treatment ^41^.

PIWIL4 cellular localization is dynamic and can shift depending on the cell’s condition. Under basal conditions, it can be found in the cytoplasm, but in response to different stimuli it shuttles between the nuclear and cytoplasmic compartments. In this study, we observed a distinct subcellular redistribution of PIWIL4 from the nucleus to the cytoplasm in response to RSV infection. A similar cytoplasmic re-localization of PIWI proteins was reported in retinal pigment epithelial (RPE) cells in response to oxidative stress following H_2_O_2_ treatment. PIWIL4 was mainly located in the nucleus at 6 hours post-treatment but became cytoplasmic as the oxidative stress continued, becoming sequestered into cytoplasmic stress granules ^42^. PIWIL4 was predominantly nuclear in cultured synovial fibroblasts from rheumatoid arthritis patients and remained nuclear in response to TNFα+IL1β/TLR-ligands ^40^. On the other hand, Jones et al. found that murine MIWI2 protein localized primarily to the cytoplasm of ciliate airway epithelial cells and that after infection with *S. pneumoniae* it remained in the cytoplasm ^11^. Ren’s group reported PIWIL4 localization both in the nucleus and cytoplasm of primary mouse choroidal endothelial cells, which was enhanced by VEGF treatment and in response to laser-induced choroidal neovascularization ^43^.

Functionally, PIWIL4 knockdown did not impair RSV replication, measured by RSV N gene expression and viral titers, indicating that PIWIL4 does not affect the viral replication machinery. In a recent study of influenza A infection in *Miwi2*-deficient mice, lack of MIWI2 led to enhanced viral clearance, although peak of viral replication was not affected ^12^. On the other hand, silencing of PIWIL4 in HIV-1 latently infected Jurkat T cells and primary CD4^+^ T lymphocytes induced reactivation of latent HIV, leading to productive viral replication in both cellular models ^44^.

Gene expression profiling revealed that PIWIL4 knockdown in response to RSV infection leads to widespread dysregulation of immune-related pathways. Notably, canonical pathway analysis identified enrichment of TNF receptor signaling and cGAS-STING pathways among upregulated genes, while downregulated genes were associated with IL-4, IL-6, IL-8, IL-12, IL-13, IL-17 along with JAK-STAT signaling. Our protein secretion data also supports that downregulation of PIWIL4 during RSV infection significantly alters virus-induced inflammatory/immune mediator expression, with changes in cytokine, chemokine and growth factor production. In a mouse model of pneumococcal pneumonia, MIWI2-deficient mice also exhibited changes in expression of inflammatory mediators, leading to increased immune cell recruitment to the lungs ^11^.

RSV infection is known to induce increased expression of mesenchymal markers (vimentin, N-cadherin), increased TGF-β, VEGF and matrix metalloproteinases (MMPs) production, expanded goblet cell populations ^45^, enhanced collagen deposition ^46^ and myofibroblast expansion ^47^, indicating EMT activation and activation of fibrosis pathways ^48, 49^. Our findings showed that knocking down PIWIL4 in RSV-infected airway epithelial cells leads to downregulation of genes associated with epithelial mesenchymal transition (EMT)/pulmonary fibrosis pathways and HIF-1α signaling, including TGF-β, VEGF, vascular endothelial growth factor receptor 1 gene (FLT1), matrix metalloproteinase-3 (MMP3), matrix metalloproteinase-9 (MMP9), fibronectin 1 (FN1), NADPH oxidase 4 (NOX4), phosphoinositide-3-kinase, regulatory subunit 5 (PIK3R5), suggesting an important role of PIWIL4 in regulating airway remodeling in response to a viral infection. These results parallel observations from Sivagurunathan’s group, who identified germline-specific PIWI-like proteins in the human retina and retinal pigment epithelium (RPE) and showed a discrete function of PIWIL4 (HIWI2) in proliferative diabetic retinopathy (PDR) ^50^. In their study, HIWI2 silencing in ARPE19 cells reduced oxidative stress-induced VEGF level and altered EMT marker expression, including E-cadherin and α-SMA. Similarly, Ren’s group found that PIWIL4 knockdown in choroidal endothelial cells, in a model of laser-induced neovascularization, led to reduced VEGF secretion ^43^. In breast cancer cells (MDA-MB-231 and MCF-7), reducing PIWIL4 levels significantly decreased components of the Transforming Growth Factor-beta (TGF-β) and Fibroblast Growth Factor (FGF) signaling pathways, impaired the cells’ ability to migrate and invade other tissues, and lowered the expression of mesenchymal markers such as N-cadherin and vimentin, consistent with a reversal of epithelial-to-mesenchymal transition (EMT) ^16^. Together, these findings highlight a role for PIWIL4 in VEGF production and EMT-related pathways across different disease contexts.

Our piRNA profiling further revealed that PIWIL4 is essential for maintaining the expression of a large subset of piRNAs in airway epithelial cells, both under control condition and during RSV infection. The loss of PIWIL4 resulted in significant up- and downregulation of hundreds of piRNAs, suggesting that PIWIL4 is required for piRNA stability or biogenesis in somatic cells. Interestingly, while many piRNAs were differentially expressed, their predicted mRNA targets showed limited overlap with the transcriptomic changes observed in PIWIL4-silenced cells. This suggests that PIWIL4 may exert its regulatory effects at least in part through a piRNA-independent mechanism. A recent study identified the function of PIWIL1 (the homolog of mouse MIWI) independent of piRNAs in pancreatic cancer metastasis. Unloaded PIWIL1 was shown to interact with the anaphase-promoting complex/cyclosome APC/C through the ubiquitin-proteasome system. This interaction activates APC/C-dependent degradation of the cell adhesion-related protein Pinin, contributing to cancer cell metastasis^51^. A piRNA-independent function of PIWIL1 was also reported in gastric cancer^52^. During mouse spermatogenesis, MIWI (mouse homologue of PIWIL1) sequesters and inhibit the activity of the ubiquitin ligase RNF8, known to play a role in histone ubiquitylation and degradation, independently of piRNAs loading ^53^.

Among the upregulated piRNAs following RSV infection, integration with the transcriptomic dataset identified several overlapping gene targets, including DNM1P47, SPATA31C2, GLB1L, ZNF652, HOXC4, ACACA, CNNM2, TNIK, and NTM. These genes are implicated in diverse cellular processes such as chromatin regulation, metabolism, vesicular trafficking, and immune modulation. For instance, ACACA encodes acetyl-CoA carboxylase alpha, a key enzyme in fatty acid biosynthesis that can influence viral replication and innate immune signaling through lipid metabolic remodeling. The upregulation of HOXC4 and ZNF652, both transcriptional regulators, may reflect an epigenetic reprogramming of epithelial cells toward a pro-survival or reparative phenotype. Additionally, *TNIK* functions as a kinase in Wnt and NF-κB pathways, linking piRNA-associated regulatory changes to inflammation and tissue remodeling processes characteristic of RSV infection.

Conversely, overlapping targets of downregulated piRNAs included DNM1P47, ERP44, GOLGA8M, IRGQ, MTAP, NISCH, NME7, PLCG1, RALGPS1, RANBP2, RAP1GAP2, and RASEF. Notably, ERP44 and GOLGA8M are involved in endoplasmic reticulum and Golgi apparatus homeostasis, suggesting that the loss of specific piRNA-mediated repression could perturb protein folding and trafficking under viral stress. PLCG1 and RAP1GAP2 are key signaling mediators in calcium and small GTPase pathways, respectively, and their deregulation may influence epithelial barrier integrity. The identification of RANBP2, a nuclear pore component with roles in interferon signaling, further underscores a potential connection between piRNA dysregulation and host defense mechanisms. Interestingly, MTAP downregulation could indicate metabolic stress or altered methylation dynamics, as this enzyme participates in the methionine salvage pathway and epigenetic regulation.

## 5 Conclusion

In conclusion, our study identifies PIWIL4 as an RSV-inducible regulator of airway epithelial cellular responses, linking it to both piRNA expression and independent gene regulatory functions, expanding our understanding of the role of PIWI proteins in somatic cells, in particular in relationship to virus-induced pathways that are critical for immune cell recruitment, epithelial barrier integrity and tissue remodeling during infection.

## RESOURCES AVAILABILITY

### Lead contact

Requests for further information and resources should be directed to and will be fulfilled by the lead contact, Tiziana Corsello.

### Materials availability

This study did not generate new unique reagents.

## SUPPLEMENTAL MATERIAL

Supplemental Excel file S1, GEN Diff genes; Supplemental Excel file S2, IPA Canonical Pathways; Supplemental Excel file S3, Diff piRNAs; Supplemental Excel file S4, Upregulated piRNAs Found target Predicted targets; Supplemental Excel file S5, Downregulated piRNAs Found target Predicted targets.

## ACKNOWLEDGMENTS

This research was supported by National Institutes of Health (NIH) grants R21 AI171664 (to A.C.), and UTMB LUDIR Pilot Grant (to A.C.); P01 AI062885 (to A.C. and R.P.G). T.C. was supported by the Institute for Human Infections and Immunity (IHII): IHII NTT; the Parker B. Francis Fellowship Program; the American Lung Association: Catalyst Award: CA-1040556. The authors would like to thank Cynthia Tribble for assistance with the manuscript tables. Figures were created with GraphPad software with an institutional license provided by UTMB.

## AUTHOR CONTRIBUTIONS

A.C. and T.C. conceived and designed research; T.C., T.L., N.D. and S.F. performed experiments; T.C., A.S.K. and Y.Z. analyzed data and prepared figures; T.C. and A.C. interpreted results of experiments and drafted manuscript; T.C., R.P.G., and A.C. edited and revised manuscript; T.C., T.L., A.S.K., Y.Z., N.D. and S.F., R.P.G., and A.C. approved final version of manuscript.

## AI DECLARATION

During the preparation of this work, the authors used Copilot tool only to improve the grammar language in certain parts of the manuscript. After using this tool/service, the authors reviewed and edited the content as needed and took full responsibility for the content of the published article.

